# What is the link between stringent response, endoribonuclease encoding Type II Toxin-Antitoxin systems and persistence?

**DOI:** 10.1101/069831

**Authors:** Bhaskar Chandra Mohan Ramisetty, Dimpy Ghosh, Maoumita Roy Chowdhury, Ramachandran Sarojini Santhosh

## Abstract

Persistence is a transient and non-inheritable tolerance to antibiotics by a small fraction of a bacterial population. One of the proposed determinants of bacterial persistence is Toxin-Antitoxin systems (TAS) which are also implicated in a wide range of stress-related phenomena. In a report (Maisonneuve E, Castro-Camargo M, Gerdes K. 2013. Cell 154:1140-1150) an interesting link between ppGpp mediated stringent response, TAS and persistence was proposed. It is proposed that accumulation of ppGpp enhances the accumulation of inorganic polyphosphate which modulates Lon protease to degrade antitoxins. The decrease in the concentration of antitoxins supposedly activated the toxin to increase in the number of persisters during antibiotic treatment. In this study, we show that inorganic polyphosphate is not required for Lon-dependent degradation of YefM, the antitoxin of YefM/YoeB TAS. The Δ10 strain, an *Escherichia coli* MG1655 derivative in which the ten TAS are deleted, is more sensitive to Ciprofloxacin and Ampicillin compared to wild-type MG1655. Furthermore, we show that the Δ10 strain has relatively lower fitness compared to the wild type and hence, we argue that the implications based on this strain are void. We conclude that there is no direct and specific link between stringent response and the regulation of TAS. The link between TAS and persistence is inconclusive due to altered fitness of Δ10 strain and hence requires thorough inspection and debate.

**Importance:** A model connecting stringent response, endoribonuclease encoding Type II Toxin-Antitoxin systems (TAS) and persistence is widely propagated. It states that “accumulation of ppGpp results in accumulation of inorganic polyphosphate which modulates Lon protease to degrade antitoxin rendering toxins free to induce persistence”. This work presents a contradiction to and challenges the model. Experimental evidence, literature survey as well as rationale are provided to show that inorganic polyphosphate is not required for the degradation of YefM, the antitoxin in YefM/YoeB TAS. The Δ10 strain is relatively more sensitive to Ciprofloxacin and Ampicillin as well as has lowered fitness. This is likely because of the polar effects on the adjacent genes caused by the genetic manipulation of multiple TAS loci.

## Introduction

Toxin–antitoxin systems (TAS) are operons consisting of two or three adjacent genes which code for a toxin, which has the potential to inhibit one or more cellular processes and an antitoxin. The antitoxin forms a complex with the toxin and suppresses the lethality of the toxin. Prokaryotic DNA sequence database mining showed that TAS are abundant in bacterial and archaeal chromosomes often in surprisingly high numbers (Anantharaman and Aravind, 2003;Pandey and Gerdes, 2005;Shao et al., 2011). Based on the antitoxin gene products, either RNA or protein, TAS are divided into 5 types (Goeders and Van Melderen, 2014) of which Type II are the most predominant and well characterized. Type II TAS encode two proteins referred to as toxin and antitoxin. They are the predominant type encoded by bacterial genomes and plasmids. The toxin has the potential to inactivate vital cellular targets while the antitoxin has the potential to sequester toxins off the cellular targets by forming a toxin-antitoxin complex (Gerdes et al., 2005;Yamaguchi and Inouye, 2011). Toxins and antitoxins also have the autoregulatory function wherein the TA complex binds to the operator present upstream of the TA operon and results in repression. The antitoxin is highly unstable and its relative concentration plays a critical role in transcriptional autoregulation as well as regulation of toxin activity (Gerdes et al., 2005;Yamaguchi and Inouye, 2011). The decrease in antitoxin concentration is a prerequisite for transcriptional activation of TAS. The significance of TAS multiplicity on prokaryotic genomes and their physiological role is highly debated (Magnuson, 2007). Many plasmids also encode TAS whose gene products have the ability to inhibit the growth of the cells cured of TA-encoding plasmids and thereby increase the population of plasmid-containing cells (Gerdes et al., 1986).

Chromosomal TAS were first discovered in studies dealing with stringent response and later in persistence. Stringent response, a response elicited in cells under amino acid starvation, is characterized by accumulation of ppGpp alarmone catalyzed by RelA upon stimulation by uncharged tRNA at the ribosomal A site (Lund and Kjeldgaard, 1972;Haseltine and Block, 1973;Cashel et al., 1996;Wendrich et al., 2002). Accumulation of ppGpp modulates RNA polymerase resulting in reduction of rRNA synthesis and thus prevents frivolous anabolism (Barker et al., 2001;Artsimovitch et al., 2004). Several mutants deficient/altered in stringent response were shown to be mutants of *relBE* (Mosteller and Kwan, 1976;Diderichsen et al., 1977), a TAS encoding an antitoxin (RelB) and a ribosome dependent endoribonuclease toxin (RelE) (Gotfredsen and Gerdes, 1998;Christensen et al., 2001). Persistence, a phenomenon of non-inheritable antibiotic tolerance, is the second instance in which genes belonging to the TA family were recognized. Some mutants, high persister mutants (*hip*), of *Escherichia coli* formed more number of persisters than the wild type. These *hip* mutations mapped to the *hipA* locus (Moyed and Bertrand, 1983) which is now recognized as a genuine TAS encoding HipA toxin and HipB antitoxin (Korch et al., 2003;Germain et al., 2013;Kaspy et al., 2013). A recent study shows an attractive link between TAS, stringent response and persistence; ppGpp, through inorganic polyphosphate (polyP), activates TAS resulting in induction of persistence (Maisonneuve et al., 2011;Maisonneuve et al., 2013). The crucial link between ppGpp and TA-mediated persistence is the essentiality of polyP for the degradation of antitoxins. During stringent response, polyP accumulates due to ppGpp-mediated inhibition of exopolyphosphatase (P_p_X) (Kuroda et al., 1997). The presence or absence of polyP determines the substrate specificity of Lon protease (Kuroda et al., 2001). Maisonneuve et al., 2013 have shown that polyP is essential for Lon-dependent degradation of YefM and RelB antitoxins resulting in increased persistence (Fig 1). YefM is the antitoxin encoded by *yefM/yoeB* TAS, a well-characterized Type II TAS. YoeB, the toxin, is a ribosome-dependent endoribonuclease (Christensen-Dalsgaard and Gerdes, 2008;Feng et al., 2013) that cleaves mRNA. YefM forms a complex with YoeB resulting in inhibition of endoribonuclease activity of YoeB (Cherny et al., 2005;Kamada and Hanaoka, 2005) and also in mediating transcriptional autorepression (Kedzierska et al., 2007).

**Figure 1.**
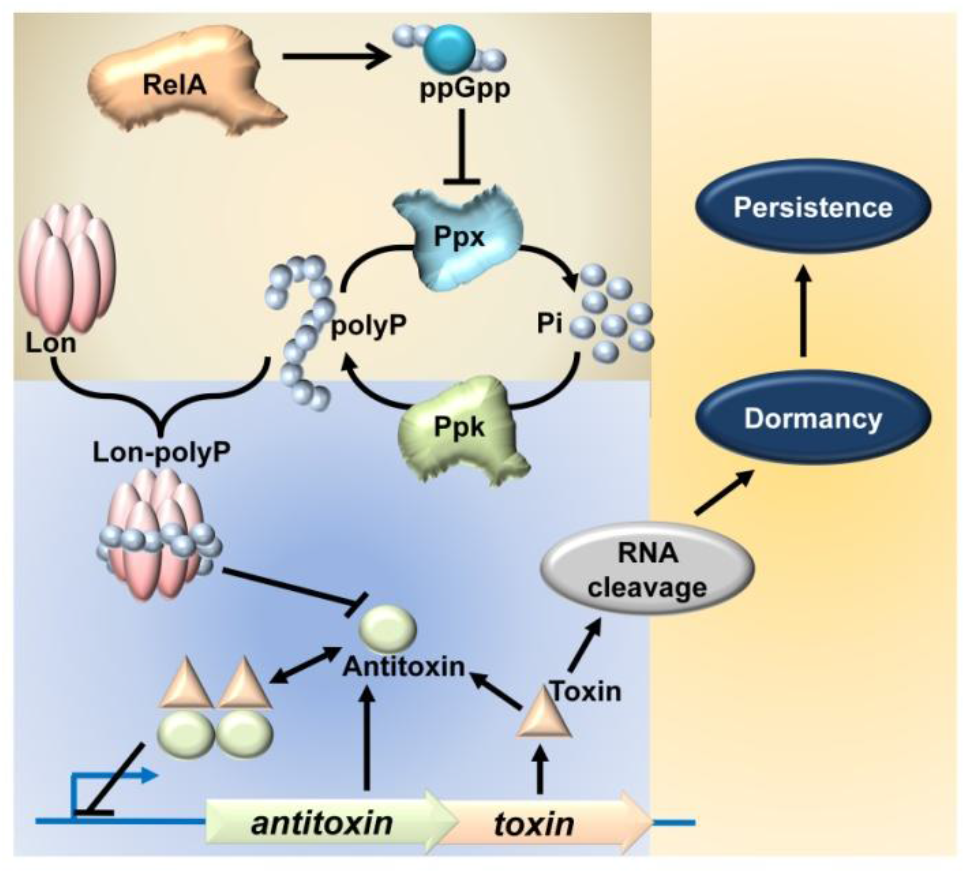
Model linking stringent response, TAS and persistence (Maisonneuve et al., 2011;Maisonneuve et al., 2013). RelA, when activated stochastically or during amino acid starvation, synthesizes the ppGpp. Accumulation of ppGpp inhibits Ppx, thereby inhibiting the degradation of polyP into inorganic phosphate. Hence, due to continual synthesis by Ppk, polyP accumulates in the cell. PolyP then modulates the substrate specificity of Lon protease, specifically targeting antitoxin proteins for degradation. This is hypothesized to render the toxin free to act on its target RNA and confer persistence.

Interestingly, in earlier studies using semi-quantitative primer extension, Prof. Kenn Gerdes’ group has concluded that transcriptional regulation of *relBE* and *mazEF* systems is independent of ppGpp (Christensen et al., 2001;Christensen et al., 2003). Hence, in this study we analyzed the essentiality of polyP in degradation of YefM antitoxin by studying the promoter activity of *yefM/yoeB* loci and endoribonuclease activity of chromosomally encoded YoeB. The endoribonuclease encoding TAS are horizontally transferring genes and most them in *E. coli* are integrated between genes with significance in bacterial stress physiology (Fiedoruk et al., 2015;Ramisetty and Santhosh, 2016). We speculated that deletion of the ten TAS could alter the physiology of the mutant strain. This study is aimed to evaluate the link between three vital aspects of bacterial stress physiology; endoribonuclease encoding TAS, stringent response, and persistence.

## Materials and methods

### Strains, plasmids and growth conditions

*Escherichia coli* MG1655, Δ5 (MG1655 derivative with 5 TAS deletions *ΔrelBE ΔmazF ΔdinJyafQ ΔyefM/yoeB ΔchpB*) (Christensen et al., 2004), MG1655 Δ*yefM/yoeB* (SC36) (Christensen et al., 2004), MG1655 Δ*relA*Δ*spoT* (Christensen et al., 2001), MG1655 Δ*ppkppx* (Kuroda et al., 1997) and Δ10 strains (MG1655 derivative with 10 TAS deletions Δ*relBE*, Δ*chpB*, Δ*mazF*, Δ*dinJ/yafQ*, Δ*yefM/yoeB,*Δ*yafNO*, Δ*hicAB*, Δ*higBA*, Δ*prlf/yhaV*, and Δ*mqsRA*). pBAD33 (Guzman et al., 1995) and pBAD-*lon* (Christensen et al., 2004) (*lon* gene cloned downstream of arabinose inducible promoter) were used for overexpression experiments. The cloning of *ppk* gene downstream of arabinose inducible promoter yielded *pBAD*-*ppk* (Mikkel Girke Jorgensen, unpublished). The cultures in all the experiments were grown in Luria Bertani broth, at 37 °C, with 180 rpm shaking in a shaker unless specified otherwise.

### Databases

EcoGene 3.0 (Zhou and Rudd, 2013) and RegulonDB (Salgado et al., 2013) were followed for nucleotide sequences, protein sequences, and regulatory information wherever required.

### Primer extension

Samples of 25 ml experimental cultures were collected at 0, 10, 30 and 60 minutes and cells were harvested by centrifugation at 4 °C. Total RNA was isolated using hot phenol method and quality was analyzed by agarose gel electrophoresis. ^32^P labelled primers, YefMPE-2 (5’-GGCTTTCATCATTGTTGCCG-3’) and lpp21 (5’-CTGAACGTCAGAAGACAGCTGATCG-3’), were used in primer extension experiments involving *yefM/yoeB* promoter activity and YoeB-dependent mRNA cleavage site mapping respectively. Reverse transcription was carried out on 10 μg of total RNA, purified from samples at designated time points, using AMV-reverse transcriptase. Sequencing reactions were carried out similarly with Sanger’s dideoxynucleotide method.

### Growth curve

12 hour old overnight cultures were grown to mid log phase in tubes in LB medium at 37 °C. The culture was rediluted 100 folds in LB medium and 2 μL of the diluted cultures were inoculated into 200 μL of LB in microtitre plate wells in triplicates. The microtitre plates were incubated at 37 °C with 170 rpm shaking. Optical density at 595 nm was measured in a 96 well microtitre plate reader (Biorad^TM^), every one hour, for 8 hours.

### Maximal CFU/ml in optimal conditions in 12 hours

Overnight cultures were inoculated into tubes containing 3 ml LB broth and grown at 37 °C with 170 rpm shaking for 12 hours. 10 μL of the culture was diluted appropriately and plated on LB plates and incubated overnight. Colonies were counted and colony forming units per ml were determined according to the dilution factor. The CFU/ml of MG1655 was taken as 100% for LB medium and culture conditions.

### Biofilm assay

Overnight cultures were diluted 100 fold in fresh LB tubes. 2 μL of inoculum was added into 200 μL of LB broth in 96 well microtitre plates. The plates were incubated at 37 °C for 16, 24, 48 and 72 hours at 37 °C. After the specified time points the plates were washed with PBS to remove floating cells. 125 μL of 1% crystal violet was added to each well and left for 20 minutes. The plates were washed with water twice and the dye was re-dissolved by adding 90% ethanol. The re-dissolved crystal violet was taken into new wells, to avoid biofilm interference, and readings were taken at 595 nm and readings were represented as the amount of biofilm formation. Experiments were carried out independently thrice in quadruplicates. Error bars indicate standard error.

### Antibiotic sensitivity test

Conventional disc diffusion method was used to measure relative sensitivity of the strains. 100μL of diluted (100 fold) overnight cultures was spread on LB agar (height – 5 mm) contained in plates with diameter 9.5 cm. Premade antibiotic discs with defined concentrations (purchased from HiMedia^TM^) were placed on the agar plates after 20 minutes. ERY = Erythromycin (15 μg), GEN = Gentamycin (10 μg), TET = Tetracycline (30 μg), NA = Naldixic acid (30 μg), AMP = Ampicillin (10 μg), CLM = Chloramphenicol (30 μg), VA = Vancomycin (10 μg), CIP = Ciprofloxacin (5 μg). The plates were incubated overnight at 37 °C. Diameters of the zones of inhibition were measured and the graph was plotted. The bars represent averages of three independent experiments done in triplicates. Error bars indicate standard error.

### Persistence assay

Exponentially growing cells of MG1655 and Δ10 were exposed to various antibiotics at the specified concentrations (Ciprofloxacin 1 μg/ml, Ampicillin 100 μg/ml, Erythromycin 100 μg/ml, Kanamycin 50 μg/ml and Chloramphenicol 100 μg/ml). After 4 hours of antibiotic treatment, cells were harvested, serially diluted and plated. After 24 hours of incubation number of viable cells was counted. Percentage of survival after antibiotic treatment for Δ10 strain is compared with the wild type MG1655 strain. The bars represent averages of three independent experiments done in triplicates. Error bars indicate standard error. AMP=Ampicillin (100 μg/ml), CIP=Ciprofloxacin (1 μg/ml), CLM=Chloramphenicol (100 μg/ml), KAN=Kanamycin (50 μg/ml), ERY=Erythromycin (100 μg/ml).

## Results and discussion

### Inorganic polyphosphate is not required for transcriptional upregulation of ***yefM/yoeB*** loci

The transcriptional upregulation of *yefM/yoeB* loci, or any typical TAS, is inversely proportional to the relative concentration of YefM. This is because TA proteins autoregulate their promoter/operator; transcription of TA operon is inversely proportaional to the concentration of antitoxin. Hence, any transcriptional activation from *yefM/yoeB* operon indicates a decrease in antitoxin concentration. Therefore, quantification of the TA mRNA is a good indicator of antitoxin concentration in the cell. To test the essentiality of polyP in Lon-dependent degradation of YefM in vivo, we employed semi-quantitative primer extension (Christensen et al., 2001;Christensen et al., 2003) of YefM mRNA. Although indirect, this assay has the advantage of a holistic transcriptional regulatory scenario of TAS without employing any genetic manipulations within the TA circuitry, thus avoiding artifacts. To test the role of ppGpp and polyP in the regulation of *yefM/yoeB* system, we performed amino acid starvation experiments using serine hydroxymate (SHX) and analyzed the transcription of *yefM/yoeB* loci using semi-quantitative primer extension using a YefM mRNA-specific primer. Exponentially growing *E. coli* strains MG1655 (Wild type), *Δlon*, *ΔppkΔppx,* and *ΔrelAΔspoT*, were treated with 1 mg/ml of SHX to induce serine starvation. *ΔppkΔppx* and *ΔrelAΔspoT* strains are deficient in accumulating polyP and ppGpp, respectively (Xiao et al., 1991;Crooke et al., 1994). In the wild type strain, we found a dramatic increase (16 fold) in the transcription of *yefM/yoeB* loci while in *Δlon* strain there was no change (Fig 2A). Interestingly and importantly, we found a higher level of transcription of *yefM/yoeB* loci in *ΔrelAΔspoT* as well as *ΔppkΔppx* strains indirectly indicates that ppGpp or polyP is not required for YefM degradation during amino acid starvation. To further investigate the essentiality of polyP we also carried out overexpression of Lon protease in MG1655 and *ΔppkΔppx* strains to know the role of polyP in the regulation of *yefM/yoeB* system and found that transcription of *yefM/yoeB* increased similarly in both MG1655 and *ΔppkΔppx* strains (Fig 2B). These observations corroborate the earlier findings that the transcriptional regulation of *relBE* (Christensen et al., 2001) and *mazEF* systems (Christensen et al., 2003) during SHX-induced starvation is independent of ppGpp but dependent on Lon protease. In fact, RelA dependent accumulation of ppGpp was shown to be inhibited by chloramphenicol treatment (Svitil et al., 1993;Boutte and Crosson, 2011) and yet the *relBE* and *mazEF* TAS were shown to be upregulated upon addition of chloramphenicol (Christensen et al., 2001;Christensen et al., 2003).

**Figure 2.**
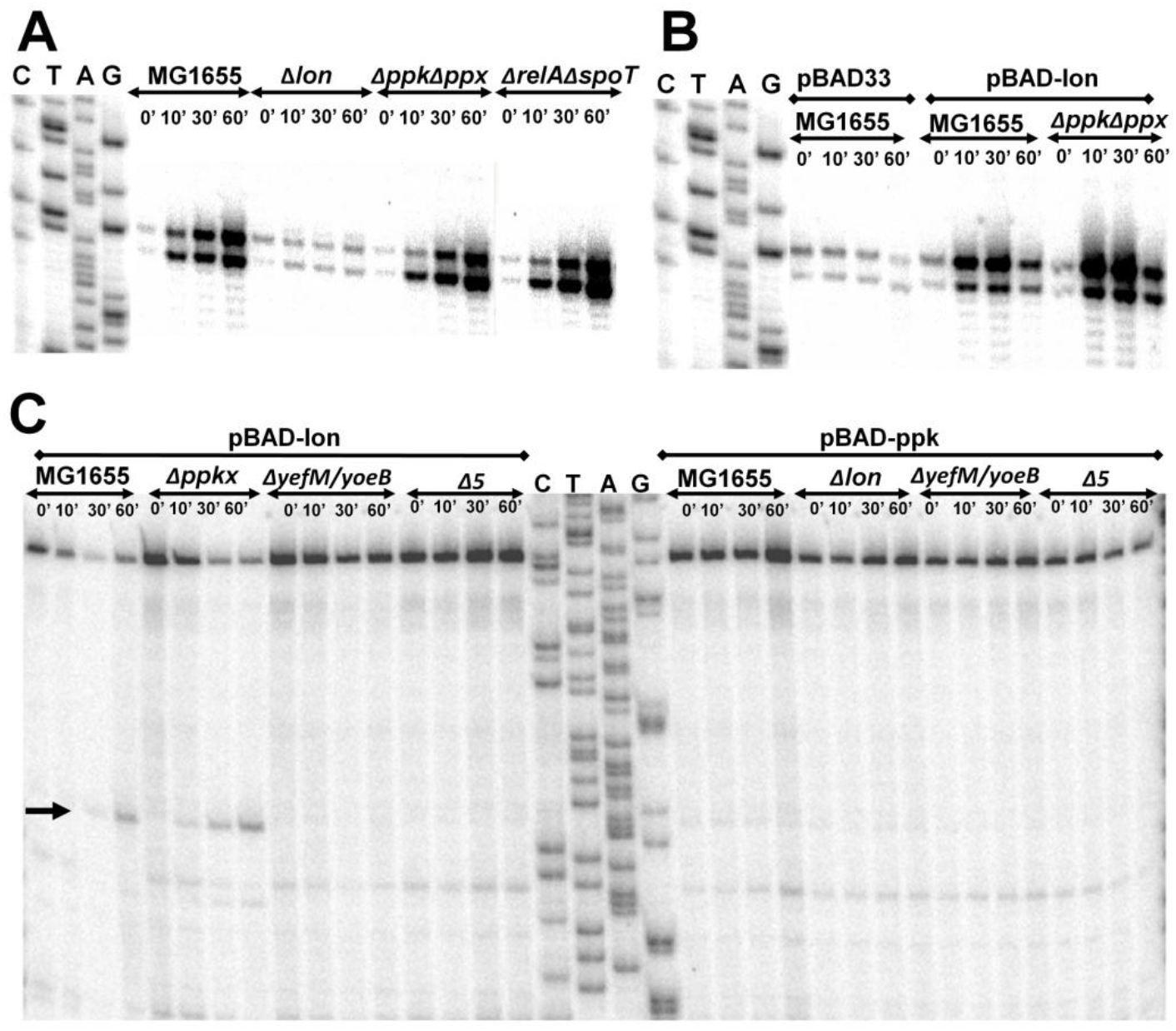
**A**. Exponentially growing (0.45 of OD_450_) cultures of MG1655, *Δlon*, *Δppkppx* and *ΔrelAΔspoT* were treated with 1 mg/ml of serine hydroxymate. Total RNA was isolated at 0, 10, 30 and 60 minutes and semi-quantitative primer extension was performed using YefM mRNA-specific primer (YefMPE-2). **B**. MG1655 and *Δppkppx* strains were transformed with pBAD-*lon* plasmids with a control of pBAD33 harbouring MG1655. Overnight cultures were diluted and grown to 0.45 OD_450_ in LB medium supplemented with glycerol as carbon source at 37 °C. Lon overexpression was induced by addition of 0.2 % arabinose. Samples were collected at indicated time intervals and semi-quantitative primer extension performed as described in Materials and methods. **C.** YoeB-dependent cleavage upon overexpression of *lon* is independent of polyP. MG1655, *ΔppkΔppx (Δppkx)*, *ΔyefM/yoeB* and *Δ5* strains were transformed with pBAD-*lon* and pBAD-*ppk* was transformed into MG1655, *Δlon*, Δ*yefM/yoeB* and *Δ5*. The transformants were grown in LB media supplemented with 2 % glycerol to mid-exponential phase (0.45 of OD_450_). 0.2 % arabinose was added to induce expression of *lon* or *ppk*. Samples were collected at 0, 10, 30 and 60 minutes and primer extension was carried out using Lpp mRNA-specific primer (lpp21) for cleavage site mapping. YoeB-dependent cleavage, indicated by an arrow, is in accordance with results from Christensen, *et al*, 2004.

Transcriptional upregulation of *yefM/yoeB* operon does not necessarily mean that YoeB is free to cleave its target mRNA. To date, chromosomal YoeB-dependent mRNA cleavage has been observed only upon ectopic overproduction of Lon protease (Christensen et al., 2004). The ectopic overexpression of Lon degrades YefM, leaving YoeB free to manifest its endoribonuclease activity. Since it was reported that Lon-mediated degradation of YefM is dependent on polyP (Maisonneuve et al., 2013), it is interesting to see if polyP is essential to render YoeB free by promoting the degradation of YefM. First, we overexpressed Lon protease in WT, *ΔppkΔppx*, *ΔyefM/yoeB* (MG1655 derivate with *yefM/yoeB* deletion) and *Δ5* (MG1655 derivate in which 5 TAS are deleted) strains and mapped for cleavage sites in Lpp mRNA by primer extension as reported in earlier studies (Christensen et al., 2004). We found that Lpp mRNA is cleaved at the second codon of AAA site in WT and *ΔppkΔppx* strains but not in Δ*yefM/yoeB* and *Δ5* strains (Fig 2C). We overexpressed *ppk* in exponentially growing cultures of wild type, *Δlon*, Δ*yefM/yoeB* and *Δ5* strains. We could not detect any YoeB-dependent cleavage of Lpp mRNA upon ectopic overexpression of *ppk* in any of the strains. This implies that YoeB-specific cleavage is independent of polyP—meaning that activation of YoeB as a result of YefM degradation is independent of polyP. Hence, our results establish that polyP is not required for the transcriptional activation of *yefM/yoeB* loci and endoribonuclease activity of YoeB which imply that polyP is not required for Lon-mediated degradation of YefM. Within the scope of the experiments, it can be argued that translation and Lon protease are the only regulators of YefM concentration. There is no experimental evidence that polyP is required to degrade all the ten antitoxins in any of the studies. Mere decrease in persister formation in Δ10 and *ΔppkΔppx* strains, but not in wild type, upon *relA* overexpression made Maisonneuve et al., 2013 to assume that degradation of the other antitoxins in *E. coli* (ChpS, DinJ, MazE, MqsA, HicB, PrlF, YafN, HigA) was also dependent on polyP (Maisonneuve et al., 2013). It is likely a fallacious assumption because “polyP-dependent TAS regulation model” (Maisonneuve et al., 2013) fails to explain how all the ten significantly divergent antitoxins of *E. coli* MG1655 could be the substrates of ‘polyP-modulated Lon’ protease. It is to be noted that YefM is degraded even in MC4100 strain (*relA1* mutant strain) (Cherny et al., 2005) which is deficient in accumulating ppGpp during amino acid starvation (Metzger et al., 1989). It may also be noted that antitoxins like YafN, HigA and MqsA (YgiT) were shown to be degraded by both Lon and Clp proteases (Christensen-Dalsgaard et al., 2010). Furthermore, based on studies on “delayed relaxed response” (Christensen and Gerdes, 2004), the half-life of RelB in MC1000 strain is approximately 15 minutes and RelB101 (A39T mutant of RelB) is less than 5 minutes. It is interesting to notice that RelB101 is degraded even in a Δ*lon* strain, indicating that some other proteases may also cleave RelB101 (Christensen and Gerdes, 2004). These literature evidences indicate that changes in primary structures of antitoxins could drastically alter their protease susceptibility and specificities. Recently, it was reported that a few TAS could induce persistence even in the absence of ppGpp (Chowdhury et al., 2016). PolyP was shown to inhibit Lon protease *in vitro* (Osbourne et al., 2014) and is reported to act as a chaperone for unfolded proteins (Kampinga, 2014) which may have significant implications in bacterial stress physiology, however, is not essential for the degradation of YefM. To our rationale, since Endoribonuclease encoding TAS propagate through horizontal gene transfer mechanisms (Ramisetty and Santhosh, 2016), minimal dependence on host genetic elements maybe preferable for TAS regulation. Within the scope of our experiments conducted in this study and based on literature, it is appropriate to state that polyP is not essential for Lon-mediated proteolysis of YefM. Although we do not have a ready explanation for this fundamental contradiction, we do not rule out His-tag interference in the proteolysis assays performed by Maisonneuve et al., 2013.

### Persistence of MG1655 and Δ10 strains to Ampicillin, Ciprofloxacin, Erythromycin, Chloramphenicol and Kanamycin

The induction of persistence (Korch and Hill, 2006;Butt et al., 2014) by over expression of toxins was challenged and shown that even proteins unrelated to toxins, during controlled over expression, induced persistence (Vazquez-Laslop et al., 2006). Maisonneuve et al., 2011 reported that Δ10 strain (*E. coli* MG1655 derivative in which ten endoribunuclease TAS were deleted) formed lesser persisters compared to wild type strain when challenged with Ciprofloxacin and Ampicillin. Since there were no reports (during the time these experiments were carried out) on the TAS mediated persistence during treatment with other antibiotics, we performed similar experiments to determine persistence to treatment of logarithmically growing cultures of MG1655 and Δ10 strains to Ciprofloxacin (1 μg/ml), Ampicillin (100 μg/ml), Erythromycin (100 μg/ml), Kanamycin (50 μg/ml) and Chloramphenicol (100 μg/ml). We found that with Ampicillin and Ciprofloxacin, Δ10 strain formed significantly lesser persisters compared to the wild type MG1655 (≈65 fold) (Fig 3A). However, we could not find significant difference in the number of persisters formed by Δ10 and MG1655 with Chloramphenicol, Erythromycin and Kanamycin. If toxin induced dormancy or metabolic regression results in bacterial persistence, it is expected that similar persistence is observed with inhibitors of translation. This is because all the toxins in the study are translational inhibitors by virtue of their endoribonuclease activity. At least with Kanamycin, since it is a bactericidal antibiotic, we expected persistence conferred by TAS. However, wild type and Δ10 strains formed equal number of persisters upon treatment with Kanamycin. Similar observation (Shan et al., 2015) with Genetamycin, another aminoglycoside antibiotic, wherein significant persistence was not observed (Wood, 2016), corroborates our findings.

**Figure 3.**
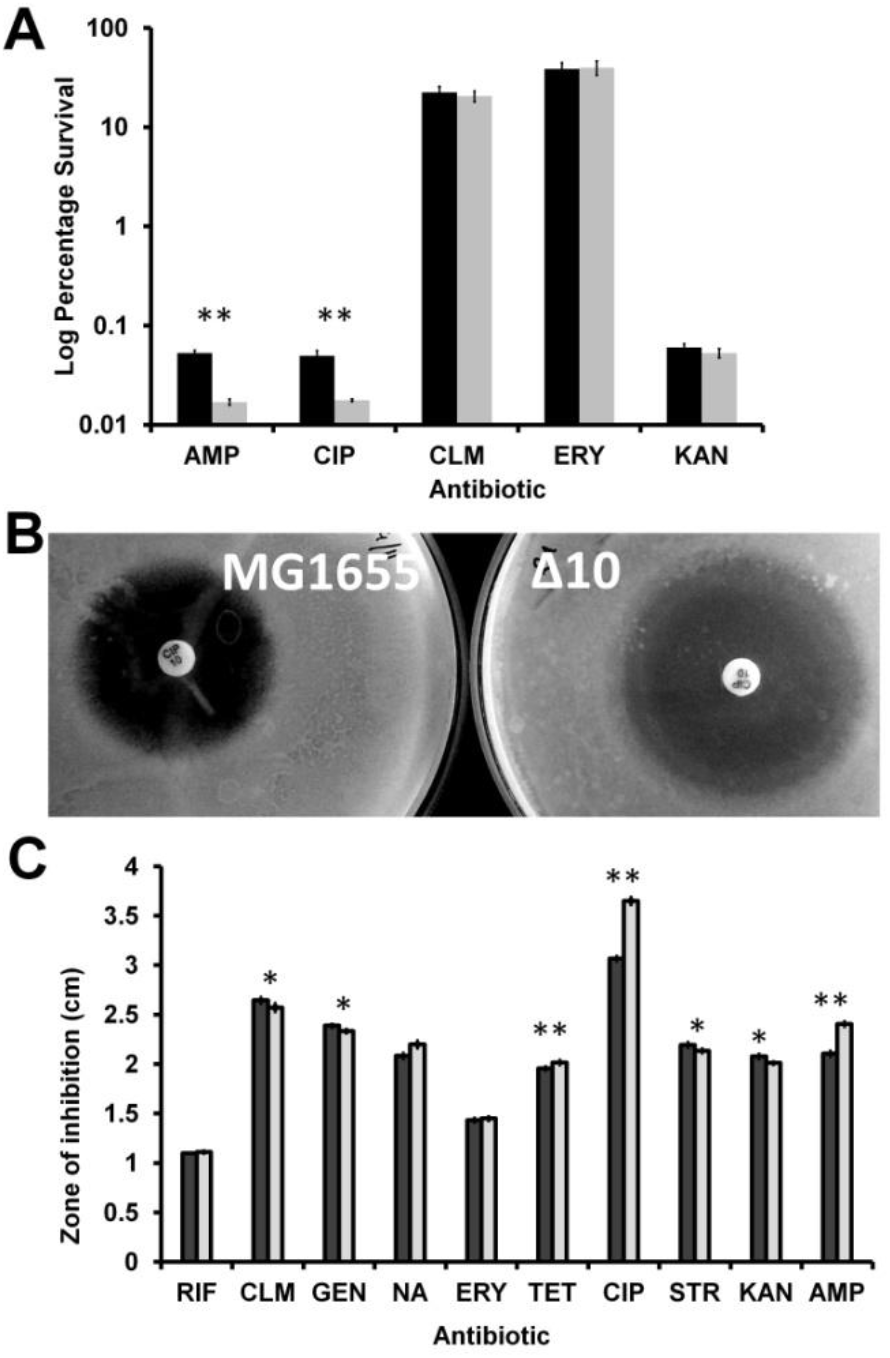
**A**. Persister cell assay with different antibiotics with mid-log phase culture. Exponentially growing cells of MG1655 and Δ10 were exposed to various antibiotics. After 4 hours of antibiotic treatment, cells were harvested, serially diluted and plated. After 24 hours of incubation number of viable cells was counted. Percentage of survival after antibiotic treatment for Δ10 strain (grey bars) is compared with the wild type MG1655 strain (solid bars). The bars represent averages of three independent experiments done in triplicates. Error bars indicate standard error. AMP=Ampicillin (100μg/ml), CIP=Ciprofloxacin (1μg/ml), CLM=Chloramphenicol (100μg/ml), KAN=Kanamycin (50μg/ml), ERY=Erythromycin (100μg/ml). (Two tailed P value ^**^ < 0.001). **B.** Overnight cultures were spread-plated on LB agar plates and antibiotic discs were placed on LA plates and incubated at 37 ^o^C for 24 hours. Representative zones of inhibition obtained for ciprofloxacin in MG1655 (left) and Δ10 (right) strains. **C.** Average diameters of the zones of inhibition, in MG1655 (black bars) and Δ10 (grey bars), around each disc loaded with premade antibiotic discs. Shown is the average of three independent experiments done in triplicates and error bars indicate Standard Deviation (SD). (P value ^**^ < 0.001, ^*^ < 0.01)

### Relative hypersensitivity of MG1655 and Δ10 strains to Ciprofloxacin and Ampicillin

In light of the above observations, we were curious about the degree of sensitivity to various antibiotics. We determined the sensitivity of the MG1655 and Δ10 strains to various antibiotics by disc diffusion method. We observed that zone of inhibition of MG1655 with Ciprofloxacin (10 μg) was 3.6 cm (averages) while that of Δ10 strain was 3 cm (Fig 3B). With Ampicillin (10 μg), the zones of inhibition for MG1655 and Δ10 strain were 2.45 cm and 2.1 cm, respectively. With Nalidixic acid, the zones of inhibition for MG1655 and Δ10 strain were 1.98 cm and 1.78 cm, respectively. We did not find any significant difference with the other antibiotics at the concentrations used. This indicates that Δ10 strain is more sensitive to Ciprofloxacin and Ampicillin relative to MG1655 (Fig. 3B&C). Our observation could also mean that TAS confers a certain degree of ‘resistance’ to antibiotics like Ciprofloxacin and Ampicillin. However, so far there are no reports that TAS confer antibiotic resistance. In light of current understanding of the role of TAS in persistence (Maisonneuve et al., 2011;Maisonneuve et al., 2013), this is an important observation. We noticed a difference in sensitivities of these strains to Ciprofloxacin and Ampicillin but not to transcription and translation inhibitors. In fact, it was reported that the Δ10 strain has lower minimal inhibitory concentration (MIC) for Ciprofloxacin (5.0 ± 0.35 ng/ml) compared to wild type (Ciprofloxacin 5.3 ± 0.45 ng/ml). Similarly, MIC of Δ10 strains was reported as 3.2 ± 0.27 μg/ml relative to MG1655 3.4 ± 0.42 μg/ml (Maisonneuve et al., 2011). Although considered as insignificant by the authors, in our view, it is irrational to infer persistence of two strains with marked difference in antibiotic sensitivities. This is vital because it deals with persistence to antibiotics and most of the experiments implicating TAS in persistence are carried out using either Ciprofloxacin or Ampicillin. It would also mean that Δ10 strain is inherently sensitive to Ciprofloxacin and Ampicillin. Although it is difficult to explain this observed sensitivity at this point of time, we do not rule out the possibility of artefacts due to genetic manipulations during the construction of Δ10 strain.

### Deletions of 10 TAS, as in Δ10 strain, causes loss of fitness

Recently we have shown that TAS are horizontally transferring genes and are integrated within the intergenic regions between important ‘core’ genomic regions (Ramisetty and Santhosh, 2016). Maisonneuve et al., 2011 reported that Δ10 strain formed lesser persisters compared to wild type strain when challenged with Ciprofloxacin and Ampicillin. Hence, we speculated that deletion of ten TAS could compromise the expression of flanking genes due to polar effects resulting in decreased fitness of the Δ10 strain. We analyzed the differences in between *E. coli* MG1655 and Δ10 strains (Maisonneuve et al., 2011) fitness by growth curve, maximal colony forming units (CFU) per ml in stationary phase and biofilm formation. We observed that the maximum growth rate (change in OD/hour) of MG1655 was 0.35 while that of Δ10 strain was 0.27 (Fig. 4A). During the 8 hour growth curve study in 96 well microtitre plates, the maximum absorbance at 595 nm was 1.05 for MG1655 while it was 0.95 for Δ10 strain (Fig. 4A). We also noticed that the optical density of the overnight cultures of Δ10 strain grown in tubes was consistently lower than that of MG1655. These observations indicated that the Δ10 strain may have metabolic deficiencies. To confirm this further we determined the CFU/ml of both the strains after 12 hours of growth in tube containing LB medium at 37 °C with 170 rpm. We observed that MG1655 yielded 7.99X10^12^ CFU/ml while the Δ10 strain yielded 4.92X10^12^ CFU/ml which is ≈40% lesser than the CFU/ml of wild type (Fig 4B). In a given set of conditions, the difference in the CFU/ml of two strains of a species is an indication of difference in their respective fitness.

**Figure 4.**
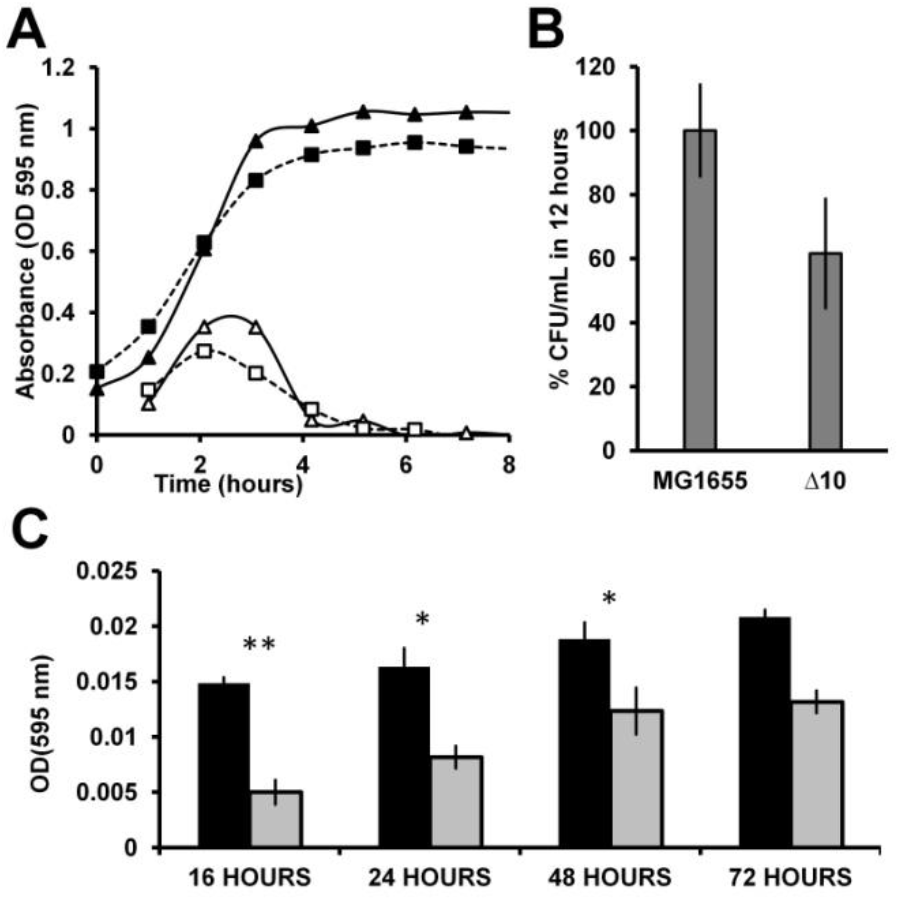
**A**. Growth curve of MG1655 and Δ10 strains. 2 μL of the diluted cultures were inoculated into 200 μL of LB in microtitre plate wells in triplicates. The microtitre plates were incubated at 37^o^C with 170 rpm shaking. Optical density at 595 nm was measured in a microtitre plate reader (Biorad^TM^). The closed triangles and closed squares represent the OD of MG1655 and the Δ10 strains respectively. The open triangles and open squares represent the growth rates (change in OD per hour) of MG1655 and the Δ10 strain respectively. **B.** Percent CFU in optimal conditions in 12 hours. Overnight cultures were incoculated into tubes containing 3 ml LB broth and growin at 37 °C with 170 rpm shaking for 12 hours. 10 μL of the culture was diluted appropriately and plated on LB plates and incubated overnight. **C.** Biofilm assay for prolonged duration. 2 μL of inoculum was added into 200 μL of LB broth in 96-well microtitre plate. The plates were incubated at 37 °C for 16, 24, 48 and 72 hours. The plates were washed and stained with1% crystal violet. Then washed thrice and the bound crystal violet was redissolved using ethanol and OD was measured at 595 nm. Experiments were carried out independently thrice in quadruplicates. Error bars indicate standard error. (Two tailed P value ** < 0.001, * < 0.01)..

We then performed biofilm assay for a prolonged period to determine any differences between these strains in their ability to form biofilms. We found that MG1655 formed consistently more biofilm, represented as absorbance of redissolved crystal violet, compared to the Δ10 strain at all the time intervals analyzed (16, 24, 48 72 hours). At 16 hours, Δ10 strain formed 66% lesser biofilm compared to MG1655 strain. Upon prolonged incubation, after 72 hours, Δ10 strain formed 35% lesser biofilm relative to wild type (Fig 4C). These observations reinforce the notion that the Δ10 strain is not as healthy as the wild type. The wild type would have the highest fitness when grown in conditions closest to their natural habitats. Mutants with one or more genetic deficiencies would lose their fitness to varying degrees. In this case, Δ10 strain has significantly lower fitness compared to the wild type likely due to the effects of deletions. The loss of fitness could be attributed to two aspects; (i) to the loss of TAS function and (ii) the polar effects on the adjacent genes due to deletion of TAS. One could argue that TAS are responsible for higher growth rate, higher CFU/ml in 12 hours as well as higher biofilm formation. However, a qualified counter argument is that the polar effects due to deletion of the ten TAS, and not necessarily the loss of TAS function, might have caused the metabolic deficiency. This is due to inadvertent interference with coding and/or regulatory sequences of the bordering regions. In our view, it is most likely that the expression of the bordering genes is compromised resulting in decreased fitness of Δ10 strain. It should be noted that TAS are horizontally transferring genes (Ramisetty and Santhosh, 2016) and are integrated within the bacterial core genome adjacent, and/or in close proximity, to important genes. As summarized in the Table 1, most of the genes that are immediately downstream of TA genes have important functions in bacterial physiology as enzymes (*yafP, fadH*) or transcriptional factors (*ydcR*, *agaR*) or in nucleotide metabolism (*mazG*, *yeeZ, ppa*) or in membrane metabolism (*yafK, hokD, ygiS*) (Table 1). It must be noted that the minimal composition of a horizontally transferring TAS consists of a promoter/operator and TA ORFs but is not composed of a terminator (Ramisetty and Santhosh, 2016). Hence, the downstream gene is cotranscribed with the TA genes because there is no promoter or terminator in the intergenic region between TA operon and the downstream gene, e.g. *relBEF* (Gotfredsen and Gerdes, 1998) and *mazEFG* (Gross et al., 2006) (Table 1). The spacers between the adjacent genes range from 9 to 218 bps, which is inclusive of the operator/promoter regions if any. Hence, the TA genes are highly linked to the downstream genes physically as well as transcriptionally. It is highly plausible that the artificial deletion of TA genes could cause polar effect on the expression of one or more of these bordering genes which is likely to result in loss of fitness (summarized in Fig. 5). Therefore, confirmation that there are no polar effects on the expression and/or the reading frames of the adjacent genes due to deletion of TA genes is essential. Attribution of the observed phenotypes solely to TAS may result in faulty interpretations and mislead the research community.

**Figure 5.**
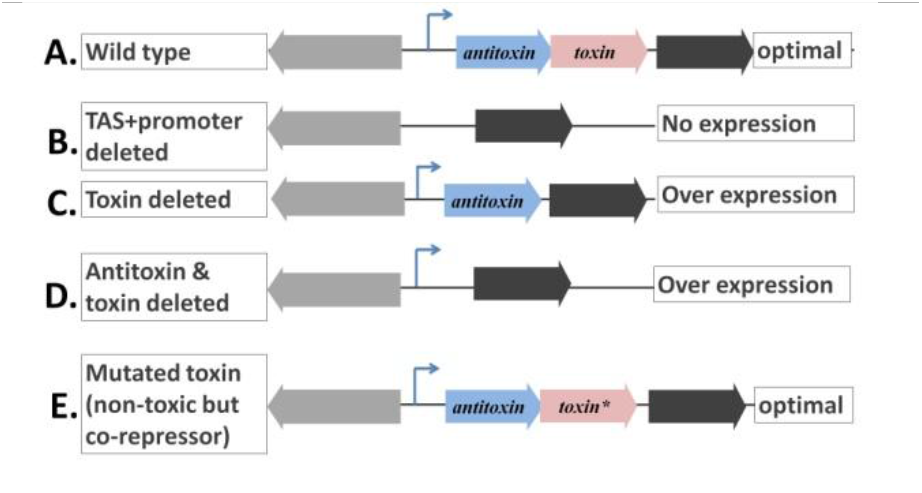
Possible polar effects on downstream gene expression due to manipulation of TAS operon. **A.** TAS operons are negatively autoregulated operons which do not encode a terminator. In the wild type, the downstream gene is cotranscribed with TA genes and its expression is considered optimal. **B.** If TAS along with promoter are deleted, there is no expression of the downstream gene due to lack of a promoter. **C.** If the toxin also is deleted, there could be over expression of the downstream gene due to missing corepressor, the toxin. **D.** If the toxin and antitoxin are deleted but not the promoter, there could be high overexpression of the downstream gene due to lack of repression. **E.** If a mutation is induced in the toxin gene such that its product is not toxic but can act as a corepressor (Overgaard et al., 2008), the expression of the downstream gene is likely to be identical to that of the wild type, an ideal case to study the function of TAS.

**Table 1.**
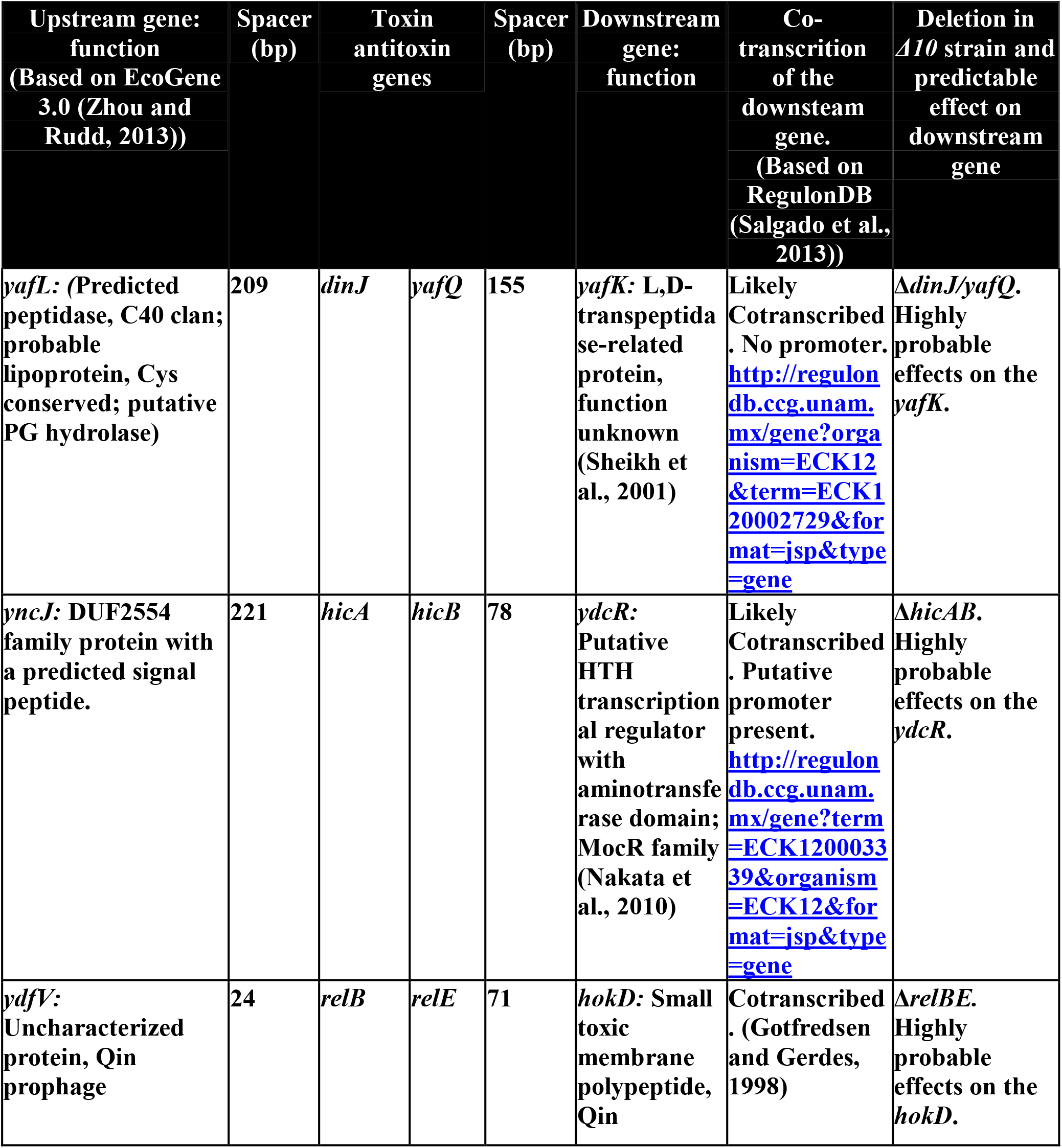
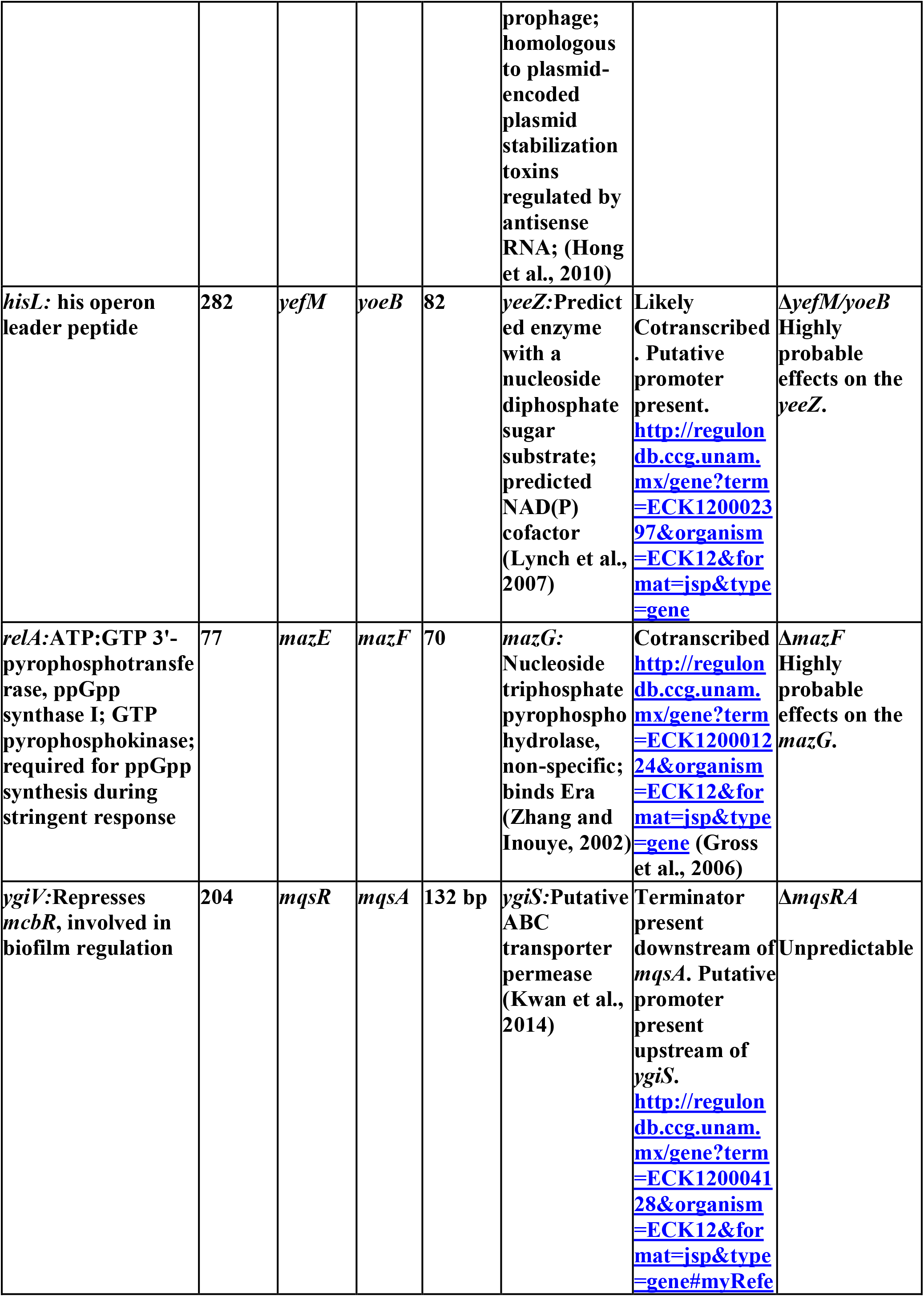
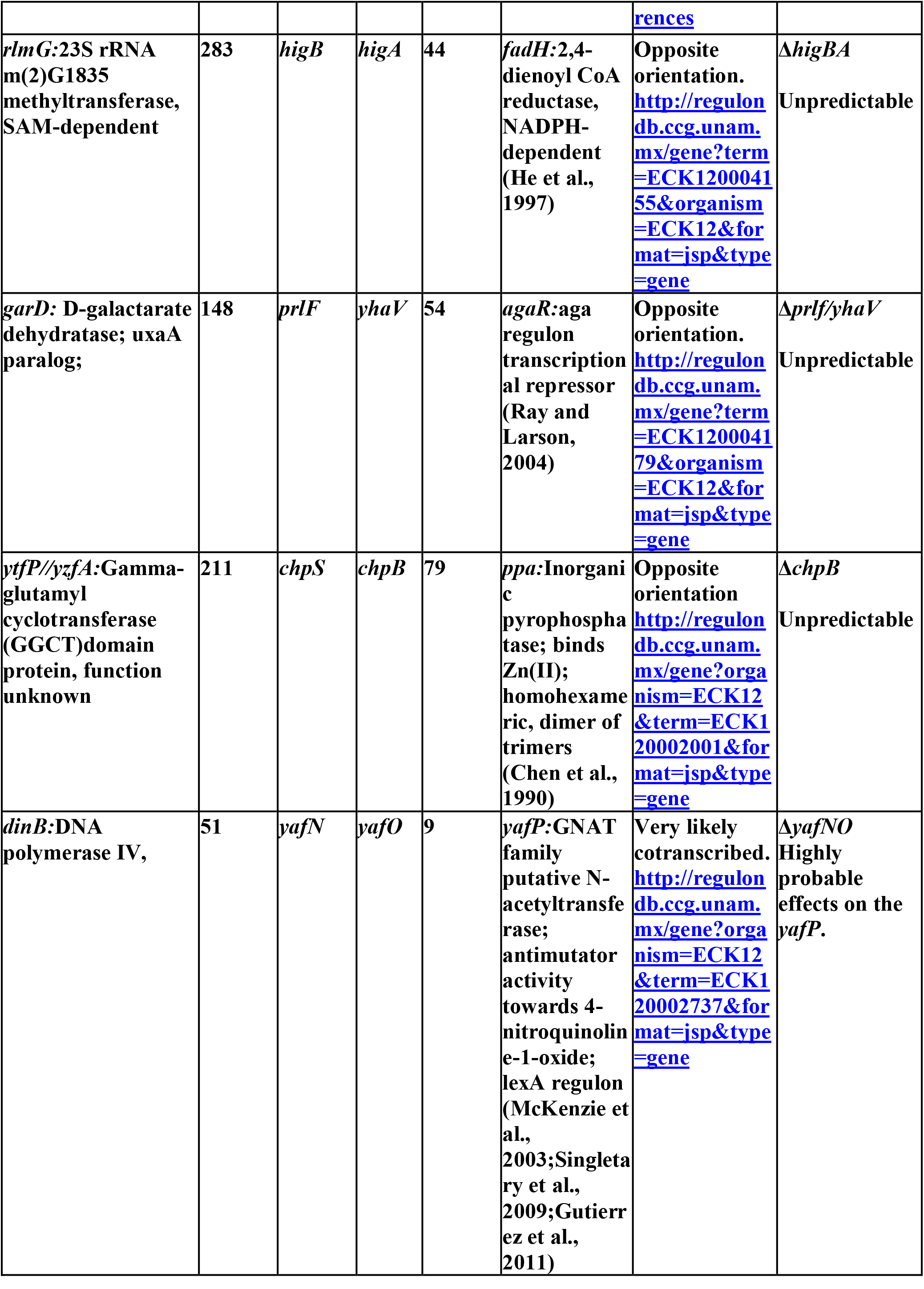
Details for the genetic organization, spacing and deletion status of each TAS in *Δ10* strain.

In the past reverse genetic studies on TAS, several deletion strains have compromised the general bacterial physiology (Gross et al., 2006;Tsilibaris et al., 2007) resulting in misleading interpretations. Construction of MC4100 *ΔmazEF* (Aizenman et al., 1996;Tsilibaris et al., 2007;Ramisetty et al., 2016) and Δ5 strain (Tsilibaris et al., 2007) strains have resulted in inadvertent interference in the coding regions of bordering genes. In a Tn-seq based genetic screen to find the molecular determinants of persisters during treatment with Gentamycin, no TAS has been found. Furthermore, in spite of having several Tn inserts in *lon* gene, *lon* mutations did not affect the persister formation frequency (Shan et al., 2015).

Toxins can induce metabolic stasis and hence we do think that TAS have the potential to induce persistence. However, more systemic studies should be carried out to definitively prove the function of TAS in persistence. As of now, with the current knowledge, we contend that chromosomal endoribonuclease encoding TAS, under their canonical autoregulatory mechanisms, may not be directly involved in persistence. We disregard ectopic overexpression of toxins’ role in persistence because it is not necessarily specific as controlled over expression of non-toxin proteins can also induce such persistence (Vazquez-Laslop et al., 2006). Similarly, we disregard the implications derived by using deletion strains such as Δ10 strain with lower fitness.

## Conclusions

The observations presented in this study, along with exhaustive literature review, establish that transcriptional control of *yefM/yoeB* is independent of polyP and ppGpp. PolyP is not required for Lon dependent degradation of YefM. Δ10 strain is relatively hypersensitive to Ciprofloxacin and Ampicillin which is probably the cause for decreased persister formation upon treatment with Ciprofloxacin and Ampicillin. Δ10 strain has lower metabolic fitness compared to wild type which also strengthens this notion. Hence the role of endoribonuclease encoding chromosomal TAS in persistence is inconclusive. Hence we refute the model presented by Maisonneuve et al., 2013. Extreme caution and evaluation should be exercised during deletion of horizontally transferring genes like TAS and evaluated for the polar effects on the downstream genes.

## Acknowledgements

We thank Prof. Kenn Gerdes for the gracious donation of the *E. coli* strains and plasmids used in this study. All primer extension work was carried out in Prof. Kenn Gerdes’ lab while at University of Southern Denmark, Odense, Denmark by BCMR. The remaining work was supported by Prof. T. R. Rajagopalan grants and Central Research Facility sanctioned by SASTRA University, Thanjavur, India.

